# Flame (v2.0): advanced integration and interpretation of functional enrichment results from multiple sources

**DOI:** 10.1101/2023.02.21.529389

**Authors:** Evangelos Karatzas, Fotis A. Baltoumas, Eleni Aplakidou, Panagiota I. Kontou, Panos Stathopoulos, Leonidas Stefanis, Pantelis G. Bagos, Georgios A. Pavlopoulos

## Abstract

Functional enrichment is the process of identifying implicated functional terms from a given input list of genes or proteins. In this article, we present Flame (v2.0), a web tool which offers a combinatorial approach through merging and visualizing results from widely-used functional enrichment applications while also allowing various flexible input options. In this version, Flame utilizes the aGOtool, g:Profiler, WebGestalt and Enrichr pipelines and presents their outputs separately or in combination following a visual analytics approach. For intuitive representations and easier interpretation, it uses interactive plots such as parameterizable networks, heatmaps, barcharts and scatter plots. Users can also: *(i)* handle multiple protein/gene lists and analyze union and intersection sets simultaneously through interactive UpSet plots, *(ii)* automatically extract genes and proteins from free text through text-mining and Named Entity Recognition (NER) techniques, *(iii)* upload single nucleotide polymorphisms (SNPs) and extract their relative genes or *(iv)* analyze multiple lists of differentially-expressed proteins/genes after selecting them interactively from a parameterizable volcano plot. Compared to the previous version of 197 supported organisms, Flame (v2.0) currently allows enrichment for 14,436 organisms.

## INTRODUCTION

Functional enrichment is the process of identifying biological functions, pathways, diseases or phenotypes where groups of genes or proteins are involved. To this end, several applications have been developed [1–5]. DAVID [6], Panther [7], Metascape [8], WebGestalt [9], Enrichr [10], aGOtool [11], AllEnricher [12], GO-Elite [13], MSigDB [14], WEGO [15], KOBAS [16], clusterProfiler [17], modEnrichr [18], agriGO [19], DOSE [20], GeneTrail [21], GOrilla [22] and ToppGene [23] are few of the widely used applications which are offered either as web tools or as software packages while others such as BiNGO [24], stringApp [25] or ClueGO [26] are offered as plugins [27] in larger frameworks such as Cytoscape [28].

While this variety of applications allows focusing on different aspects such as enriching for *(i)* biological pathways (e.g., KEGG [29], WikiPathways [30], Reactome [31]), *(ii)* gene ontologies (e.g., GO [32]) such as biological processes, molecular functions and cellular components, *(iii)* diseases (e.g., OMIM [33], DisGeNet [34]), *(iv)* protein complexes (e.g., CORUM [35]), *(v)* protein domains (e.g., Pfam [36]), *(vi)* phenotypes (e.g., HPO [37]) or *(vii)* regulatory motifs (e.g., TRANSFAC [38], miRTarBase [39]), an obvious selection of the right tool to cover certain needs is most of the times not straightforward. Moreover, while most of these tools come with a simple interface, little emphasis has been given on the visualization and combination of results, id conversion, parameter selection, result prioritization, record association, support of multiple lists and versatile input options. Therefore, usage and interpretation of results in a more streamlined and combinatorial aspect still remain difficult tasks for the average user.

In this article, we present Flame (v2.0), an advanced service for addressing the aforementioned issues. Flame focuses on usage simplicity, offering many degrees of freedom regarding importing and handling multiple lists, running advanced state-of-the-art functional enrichment analysis pipelines as well as combining and prioritizing results through advanced interactive visualizations. Flame is able to report results as lists but also associate them at a network level [40], while it offers major integration of resources [41] such as aGOtool, g:Profiler, WebGestalt, Enrichr and STRING [42]; thus referring to their broad audience while taking full advantage of their strengths.

## METHODS AND RESULTS

### Input options

Compared to its predecessor [43], Flame (v2.0) comes with a variety of newly introduced input options (Figure 1). These are mainly: *(i)* support of simple gene lists, *(ii)* support of variants and polymorphisms, *(iii)* support of text-mining and Named Entity Recognition (NER) techniques to process free text and mine it for genes and proteins and *(iv)* support of gene expression data derived from experiments.

**Figure 1.**
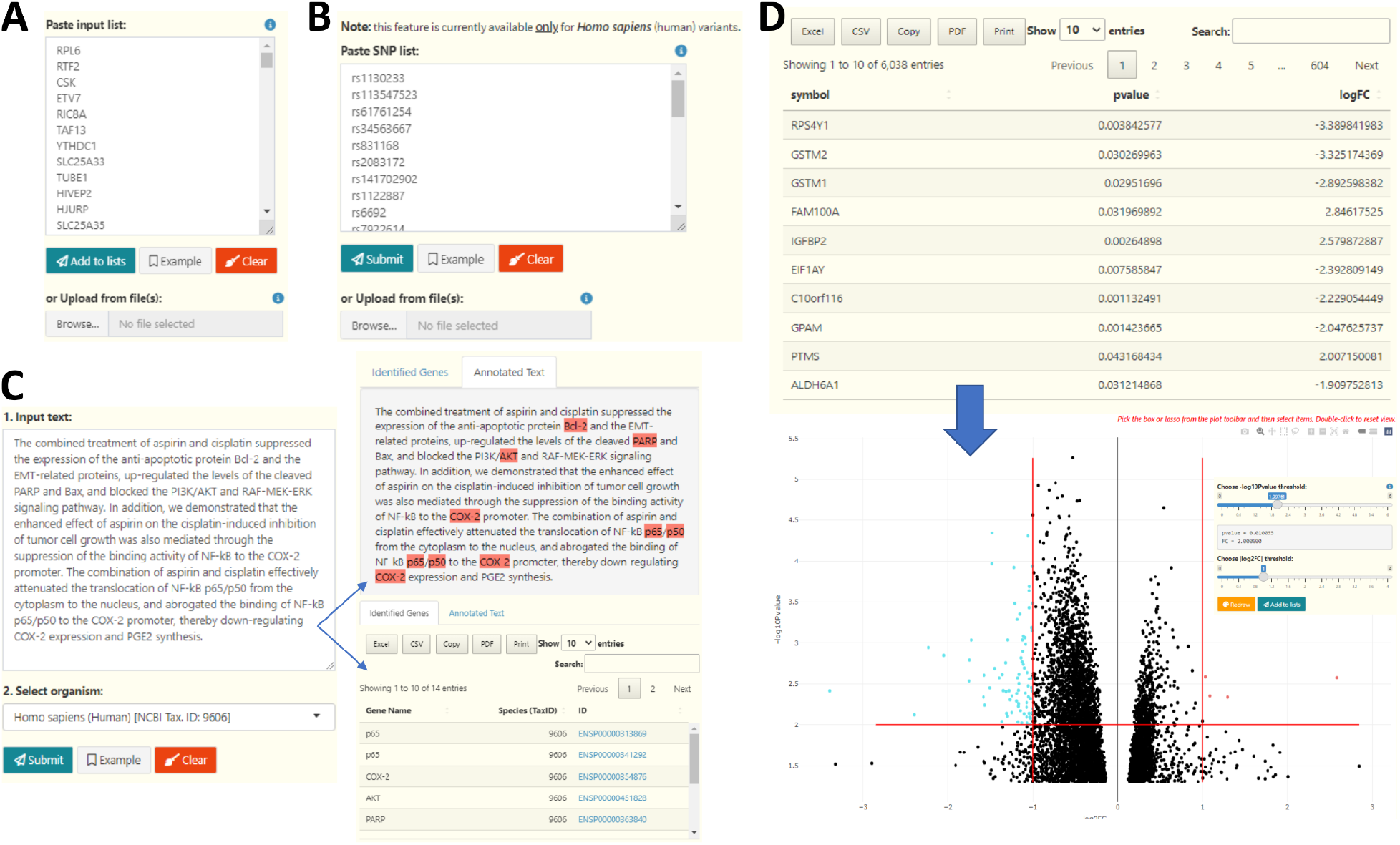
Flame input options. A) A simple gene list using any of the widely-used identifiers (e.g., HUGO names, ENSEMBL, Entrez, Uniprot). B) A list with Human SNPs. C) A free text which can be further mined for genes and proteins using NER for an organism of preference. The identified genes and proteins can be then used for functional enrichment analysis. D) Input of gene expression data (gene name, *p*-value, logFC) to generate a Volcano plot. Users can manually apply thresholds on the *p*-value and the FC axes to define the differentially over- and under-expressed genes of significance and interactively select them from the plot for further analysis.

### Gene lists

Similarly to Flame v1.0, users can directly paste or upload multiple lists by using a plethora of gene identifiers or chromosomal intervals. Gene and protein names can be imported in the Ensembl [44], Entrez [45], Uniprot [46], RefSeq [47], EMBL [48], ChEMBL [49] and WikiGene [50] namespaces, or be later converted into another namespace during the functional enrichment analysis procedure.

### Variants and Polymorphisms

In addition to directly uploading gene lists, Flame is capable of parsing and annotating lists of single nucleotide polymorphisms (SNPs), through the “SNP” input option. The input in this case is a list (whitespace or comma-delimited) of SNP rs-codes in the dbSNP [51] format. The ids are mapped to their corresponding genes and associated Ensembl identifiers through the g:SNPense functionality of g:Profiler and are then searched against dbSNP to retrieve additional metadata, including the chromosome number, strand, genomic coordinates and the variant’s reported effects. The annotated SNPs can be downloaded, while their associated genes can be added to the list of Flame inputs, using either their Entrez gene name or their Ensembl ids. Since dbSNP has dropped support for non-human variants in recent versions, this functionality is currently available only for *Homo sapiens* (human).

### Text-Mining and NER

In this version, users can provide free text and utilize the EXTRACT [52] tagger to perform Named Entity Recognition (NER) to mine it for genes and proteins for an organism of preference. Upon completion, Flame will report the identified genes/proteins in an interactive and easy-to-filter table as well as an annotated text with the identified terms highlighted. In mouse-hover, a popup window will appear, providing information about the entity as well as direct links to the STRING and Ensembl databases. The annotated terms can be added to the list of Flame inputs, using their associated Ensembl ids. This functionality is supported for all currently available organisms in Flame (∼14,000 taxa). To this end, it is worth mentioning that the EXTRACT tagger is already being used by a plethora of established text-mining based applications [42,53–55].

### Gene Expression data

Defining the significant differentially over- and under-expressed genes is most times an empirical task performed by experimentalists, who usually apply thresholds within an accepted range for parameters such as the fold change (e.g., absolute FC > 2) and the statistical significance (e.g., *p*-value < 0.05). Therefore, slightly tuning the parameters may result in a more complete biological story. For this purpose, Flame offers the ability to upload expression data in tab delimited or comma separated format (gene name - log fold change - statistical significance) and interactively select sets of genes on a generated Volcano plot. Users can adjust both the *p*-value and FC thresholds and redraw the plot, highlighting the overexpressed genes in red and the under-expressed genes in blue. Following, users may apply a selection lasso or rectangle to mark gene sets of interest directly on the Volcano plot, and append them to the Flame lists which can then be further processed and enriched.

### Integration and functional enrichment for multiple organisms from various sources

Due to the lack of standards in the field [4], functional enrichment applications often differ on the databases they support, the way of processing data, the organisms they cover, their ranking and scoring schemes, their reporting and visualizations as well as the input options they offer. While such differences may hinder usage, a combination of such tools as a means of complementing each other’s weaknesses is always an option. However, familiarity with each tool of choice is necessary but often a time consuming process. In Flame (v2.0), we overcome this issue by bringing together four established applications, namely *aGOtool, gProfiler, WebGestalt* and *enrichR* under the same interface and making them available Variants, in a consistent and unified manner.

In detail, aGOtool (and subsequently Flame) currently supports 14,436 organisms corresponding to UniProt reference proteomes, including animals (mammals, insects, fish etc.), plants, fungi, protozoa and bacteria. The tool offers functional enrichment for biological processes, molecular functions and cellular components through Gene Ontology, pathway enrichment through KEGG, Reactome and WikiPathways, disease enrichment through the Disease Ontology [56], annotation enrichment for proteins through Uniprot, Pfam and InterPro [57] as well as tissue enrichment through the Brenda Tissue Ontology [58]. It also offers literature enrichment through PubMed. Regardless of the input identifiers, gene/protein names are converted to the Ensembl protein namespace (ENSP) through the STRING API before the enrichment process. The ENSP namespace returns the most accurate results for the aGOtool enrichment pipeline.

While using the g:Profiler pipeline, extra features such as enriching for tissues through the Human protein Atlas (HPA) [59], protein annotations through CORUM, phenotypes through the Human Phenotype Ontology (HPO) [37], or DNA regulatory motifs through TRANSFAC and miRTarBase are made available. It is worth mentioning that g:Profiler comes with an advanced id conversion function which enables namespace and orthology conversion for 821 organisms, while it provides great flexibility on the identifiers which it accepts as input.

When opting for WebGestalt, users can further enrich their lists for OMIM, DisGeNet and GLAD4U [60] diseases, PANTHER pathways [7] as well as DrugBank and GLAD4U drug annotations [61] while genes/proteins are automatically converted to Entrez accession identifiers. Currently WebGestalt supports twelve organisms, namely, *Homo sapiens* (human), *Mus musculus* (mouse), *Rattus norvegicus* (rat), *Bos taurus* (bovine), *Arabidopsis thaliana* (thale cress), *Drosophila melanogaster* (fruit fly), *Caenorhabditis elegans* (nematode worm), *Saccharomyces cerevisiae* (yeast), *Danio rerio* (zebrafish), *Sus scrofa* (pig), *Canis lupus familiaris* (dog) and *Gallus gallus* (turkey).

Finally, the enrichR library offers functional enrichment for seven organisms, namely *Homo sapiens* (human), *Mus musculus* (mouse), *Bos taurus* (bovine), *Drosophila melanogaster* (fruit fly), *Caenorhabditis elegans* (nematode worm), *Saccharomyces cerevisiae* (yeast) and *Danio rerio* (zebrafish), preferably in the Entrez gene namespace and supports extra enrichment options for phenotypes through the KOMP2 Mouse Phenotypes, WormBase [62] and MGI Mammalian Phenotype [63] databases.

A summary and a direct comparison of the supported features, cutoff types and allowed namespaces are shown in Table 1. Regarding execution times, aGOtool is the default selected enrichment tool of Flame and performs the fastest while WebGestalt’s API executes the slowest and may affect Flame’s performance.

**Table 1.** Comparison of the four different enrichment tools integrated in Flame.

|  | **aGOTool** | **g:Profiler** | **WebGestalt** | **enrichR** |
| --- | --- | --- | --- | --- |
| **Organisms** | 14,436 | 821 | 12 | 7 |
| **Namespaces** | Ensembl IDs,<br>UniProt ACs,<br>STRING IDs | Ensembl, Entrez,<br>Uniprot, RefSeq,<br>EMBL, ChEMBL,<br>WikiGene | Entrez gene<br>accession<br>identifiers | Entrez gene<br>names |
| **Significance**<br>**metrics** | *p*-value, False<br>Discovery Rate<br>(FDR) | *g:SCS* (adjusted<br>*p*-value), FDR,<br>Bonferroni | Bonferroni & five<br>FDR types (BH, BY,<br>Holm, Hochberg,<br>Hommel) | Adjusted *p*-value |
| **Supported databases** |  |  |  |  |
| **Gene Ontology** | Biological Processes, Molecular Functions, Cellular Components | Biological Processes, Molecular Functions, Cellular Components | Biological Processes, Molecular Functions, Cellular Components | Biological Processes, Molecular Functions, Cellular Components |
| **Biological Pathways** | KEGG, REACTOME, WikiPathways | KEGG, REACTOME, WikiPathways | KEGG, REACTOME, WikiPathways, PANTHER | KEGG, REACTOME, WikiPathways, PANTHER |
| **Diseases** | Disease Ontology | × | OMIM, DisGeNET, GLAD4U | Disease Ontology |
| **Drugs** | × | × | DrugBank, GLAD4U | × |
| **Protein families, domains & complexes** | InterPro, Pfam, UniProt | CORUM | × | × |
| **Tissues** | BRENDA Tissue Ontology | Human Protein Atlas | × | WormBase |
| **Phenotypes** | × | Human Phenotype Ontology (HPO) | Human Phenotype Ontology (HPO) | Human Phenotype Ontology (HPO), KOMP2 Mouse Phenotypes, WormBase, MGI Mammalian Phenotype |
| **Regulatory DNA motifs** | × | TRANSFAC, miRTarBase | × | × |

### Visualization options

Flame extends its functionality by offering a plethora of visualizations and network analysis options for the interpretation of enrichment results. Besides simple searchable and interactive tables, it offers several visual alternatives for presenting the reported results from each enrichment tool (Figure 2). Flame provides four different main visualization options to represent the top enriched terms. These are: *(i)* networks, *(ii)* heatmaps, *(iii)* barcharts and *(i)* scatter plots. For each category, one can adjust the top hits interactively by setting up filters and parameters such as datasources, number of visualized top terms and scoring metric. In the case of networks and heatmaps, three different modes of association are offered, namely *Functions Vs Genes, Functions Vs Functions* and *Genes Vs Genes*. In the first mode, the top functions and their associated genes are presented either as a network (nodes linked with edges), or as a heatmap (associations are shown as colored cells/blocks). In the second mode, functions are connected with other functions based on the number of their shared genes, whereas in the third mode, genes are connected based on the common functions they are involved in. In the cases of networks, bar charts and scatter plots, one can adjust the top hits by combining multiple types of resources (e.g., pathways from KEGG, Reactome and WikiPathways). In addition, while most of the network associations are shown in 2D, Flame has the option to call Arena3D^web^ [64,65] in order to visualize heterogeneous information in a 3D view as a multi-layered graph (each biomedical entity type corresponding to a different layer). All aforementioned plots are also available for the literature enrichment pipeline, where instead of functions, PubMed articles are returned. A Manhattan plot can also be generated for g:Profiler exclusively. Finally, as an extra analysis step, Flame utilizes the STRING API to generate protein-protein interaction networks.

**Figure 2.**
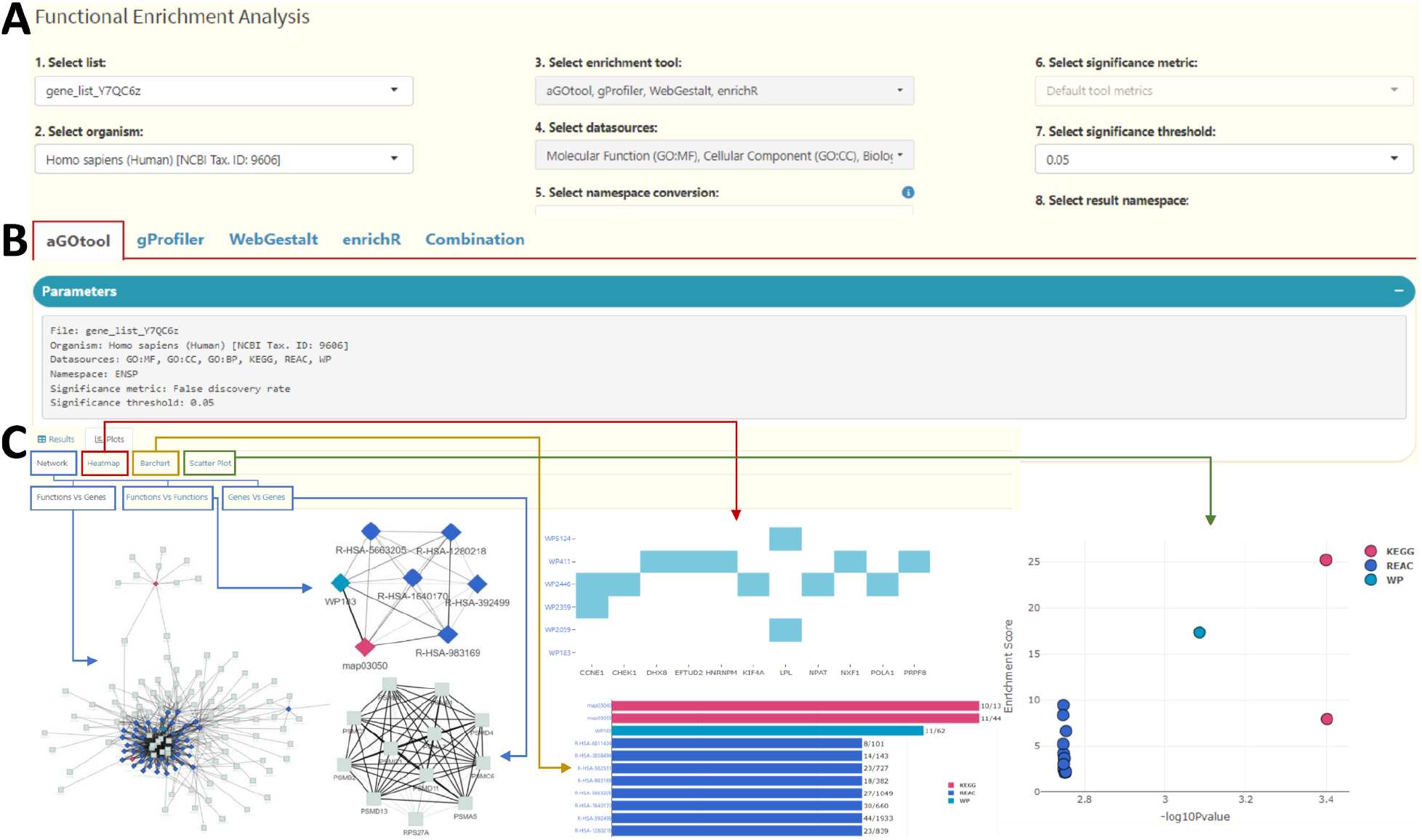
Summary of Flame’s functionality. A) Selection of list, organism, enrichment tool(s), datasources, namespace, significance metric and threshold. B) aGOtool, g:Profiler, WebGestalt and enrichR results are reported on separate tabs as well as unified into a tab called “Combination”. C) Visualization options for reporting the top enriched terms (networks, heatmaps, bar charts and scatter plots). Networks and heatmaps can be used to show *function-gene, function-function* and *gene-gene* associations while top hits from multiple sources can be shown in combination as a multi-type network, bar chart or scatter plot.

### Use of UpSet plots for comparing input lists and reported results from various sources

Flame (v1.0) uses UpSet plots, as a replacement for Venn diagrams, in order to show unions, intersections and distinct combinations among the input gene lists. Following the same approach, in this version, Flame (v2.0) extends this functionality to show how the four supported back-end pipelines (*aGOtool, gProfiler, WebGestalt, enrichR*) agree on the results they report (Figure 3).

**Figure 3.**
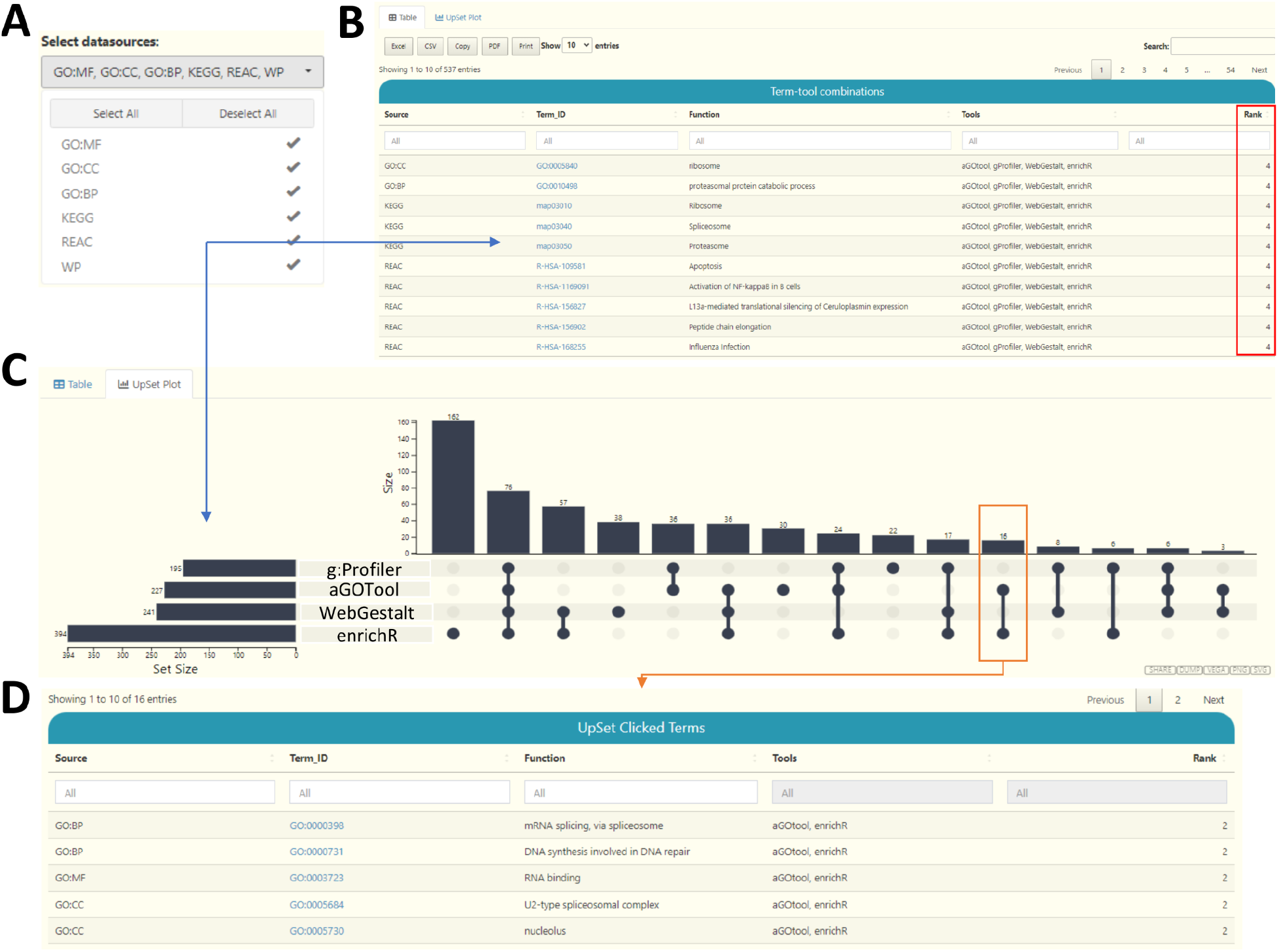
Distinct intersections of reported results. A) A selection box to limit the comparisons to certain biomedical entities. B) A searchable and interactive table reporting the intersecting biomedical entries from each tool. C) A visual representation of the distinct intersections with the use of interactive Upset plots. D) Part of a table showing 5 of the 16 biomedical entries of the distinct intersection set which corresponds to the chosen UpSet bar.

To this end, when more than one functional enrichment tool has been executed, an extra tab named “*Combination*” is generated automatically. In this tab, distinct intersections between the four resources are summarized in a searchable and easy-to-filter table as well as in an interactive UpSet plot. The table reports the aggregated enriched terms along with the tool(s) that generated the term and its respective rank (number of tools that returned it). Rank 1 means that a biomedical entity (e.g., an enriched pathway) was detected by one resource only, while rank 4 means that all four tools fetched the same result. In parallel, Flame (v2.0) generates an interactive UpSet plot where users can click on a bar to generate a table which corresponds to the subset of shared terms shown in the corresponding UpSet column. In this implementation, the UpSet plot consists of two to four rows (one for each enrichment tool) while columns describe the distinct intersections among results. Both the table and the UpSet plot can be limited to a user-selected set of biomedical entries (e.g., compare reported pathways, or biological processes or both).

### Implementation

Flame is mainly written in R (Shiny framework). The enrichment analysis with aGOtool is executed through its corresponding API (https://agotool.org/api_orig). Enrichment analysis with g:Profiler, WebGestalt and Enrichr is executed through their R libraries; *gprofiler2, WebGestaltR* and *enrichR* respectively. Enriched term networks are generated through the *igraph* R package and visualized via the *visNetwork* R library. The volcano plot input along with all other enriched term plots (heatmaps, barcharts and scatter plots) are generated with the use of the *plotly* R library. A Manhattan plot is also offered, specifically for the g:Profiler results, through its respective package. Finally, UpSet plots are drawn with the use of *upsetjs* R library.

The SNP input and namespace/orthology conversions are handled through the g:Profiler package. Text-mining input is handled by the EXTRACT tagger API (https://tagger.jensenlab.org). The PPI network analysis is offered by the STRING API (https://string-db.org/api). Flame handles its GET API requests through R and its POST API requests through a dedicated node.js server.

### Application Programming Interface

Flame comes with its own API and currently supports both GET and POST requests. Shorter requests can be done with a GET request using the following format:

https://bib.fleming.gr:8084/app/flame/?url_genes=MCL1,TTR;APOE,ACE2;TLR4,HMOX1

In this example, gene lists are separated with semicolon (;) and genes with comma (,). Longer requests can be performed via Flame’s POST request at:

https://bib.fleming.gr/bib/api/flame

using the following JSON object format:

`{“gene_list1”: [“GNAS”, “ABCG2”, “WT1”, “CDK2”, “FLT1”, “HBA1”, “CCN2”, “MDM2”],`

`“gene_list2”: [“MMP1”, “PTGS2”, “PON1”, “LDLR”, “HBA1”, “CYP1B1”, “PTEN”, “SNCA”],`

`“gene_list3”: [“UGT1A1”, “CDH1”, “MDM2”, “EGFR”, “FMR1”, “VEGFA”, “ERCC1”]}`

## DISCUSSION

Flame (v2.0) expands on its previous capabilities in many aspects (Table 2); from offering users a plethora of different input options, to a significant organism-space expansion, the addition of multiple enrichment libraries/APIs and the combination of their results, as well as through enhancing its own API. We believe that the multiple input options along with STRING’s organisms, make Flame a very competitive application which greatly enhances user experience. By also offering the streamlined execution of multiple functional enrichment tools and the aggregation of their results, we facilitate the knowledge extraction of analysis, and partially tackle the question “which tool should I use/trust?”. Given its accessibility and ease of use, we hope Flame will quickly become the go-to tool for advanced functional enrichment analysis and visualization.

**Table 2.** Main differences between Flame versions.

|  | Flame (v1.0) | Flame (v2.0) |
| --- | --- | --- |
| **Enrichment Categories** | 1. gene ontology<br>2. biological pathways<br>3. proteins<br>4. phenotypes<br>5. tissues<br>6. regulatory motifs<br>7. literature enrichment | 1. gene ontology<br>2. biological pathways<br>3. proteins<br>4. phenotypes<br>5. tissues<br>6. regulatory motifs<br>7. literature enrichment<br>8. drug associations<br>9. protein families<br>10. diseases |
| **Enrichment Tools** | 1. g:Profiler<br>2. aGOtool (only for literature enrichment) | 1. aGOtool<br>2. g:Profiler<br>3. WebGestalt<br>4. Enrichr |
| **Supported Organisms** | 197 | 14,436 |
| **Input Options** | 1. gene/identifier lists | 1. gene/identifier lists<br>2. SNP lists<br>3. NER and text-mining on a free text<br>4. gene selection from an interactive volcano plot |
| **List comparison with the use of UpsetPlots** | 1. input gene lists | 1. input gene lists<br>2. reported results from any of the four supported tools |
| **Other types of analysis** | 1. Protein-protein interaction networks<br>2. Gene orthology search<br>3. Database id conversion | 1. Protein-protein interaction networks<br>2. Gene orthology search<br>3. Database id conversion<br>4. Mapping of SNPs to genes |
| **Visualization** | 1. Plots (Bar, UpSet, Scatter, Manhattan)<br>2. Heatmaps<br>3. 2D Networks with the use of VisNetwork | 1. Plots (Bar, UpSet, Scatter, Manhattan)<br>2. Heatmaps<br>3. 2D Networks with the use of VisNetwork<br>4. 3D multi layer graphs with Arena3D<sup>web</sup> |
| **API requests** | 1. GET | 1. GET<br>2. POST |

## AVAILABILITY

**Web Application**: http://flame.pavlopouloslab.info

**Code:** https://github.com/PavlopoulosLab/flame

**Docker:** https://hub.docker.com/r/pavlopouloslab/flame

## Acknowledgements

F.A.B was supported by Fondation Santé and the Onassis Foundation. E.K was supported by the Hellenic Foundation for Research and Innovation (H.F.R.I) under the ‘First Call for H.F.R.I Research Projects to support faculty members and researchers and the procurement of high-cost research equipment grant’, Grant ID: 1855-BOLOGNA. G.A.P. was supported by the project ‘The Greek Research Infrastructure for Personalized Medicine (pMedGR)’ (MIS 5002802), which is implemented under the Action ‘Reinforcement of the Research and Innovation Infrastructure’, funded by the Operational Program ‘Competitiveness, Entrepreneurship and Innovation’ (NSRF 2014-2020) and co-financed by Greece and the European Union (European Regional Development Fund). P.K. and P.B. were supported by the project “GENOMIC OASIS: GENOMIC Analysis of Organisms of Agricultural and liveStock Interest in Sterea”, grant number 5045902. PS was supported by the Onassis Foundation.

